# TurboID-mediated proximity labelling of cytoophidium proteome in *Drosophila*

**DOI:** 10.1101/848283

**Authors:** Bo Zhang, Yuanbing Zhang, Ji-Long Liu

## Abstract

Proximity-based biotinylation combined with mass spectrometry has emerged as a powerful approach to study protein interaction networks and protein subcellular compartmentation. However, low kinetics and the requirement of toxic chemicals limit the broad utilisation of current proximity labelling methods in living organisms. TurboID, the newly engineered promiscuous ligase, has been reported to label bait proteins effectively in various species. Here, we systematically demonstrated the application of TurboID-mediated biotinylation in a wide range of developmental stages and tissues, and we also verified the feasibility of TurboID-mediated labelling in desired cells via cell-type-specific GAL4 driver in *Drosophila*. Furthermore, using TurboID-mediated biotinylation coupled with mass spectrometry, we characterized the proximate proteome of the cytoophidium, a newly identified filamentous structure containing the metabolic enzyme CTP synthase (CTPS) in *Drosophila*. Our study demonstrates a referable tool and resource for research in subcellular compartments of metabolic enzymes in vivo.

## Introduction

Affinity purification coupled with mass spectrometry (AP-MS) has been a powerful approach to identify the interaction network for organelles or protein complexes (Dunham et al., 2012; Gingras and Raught, 2012). However, AP-MS shares some important limitations in the detection of weak/transient interactions, especially in insoluble subcellular compartments, such as membrane, chromatin, nuclear lamina or the cytoskeleton (Gingras et al., 2019; Lambert et al., 2015; Qi and Katagiri, 2009; Van Leene et al., 2007).

The recent development of proximity-dependent labelling methods has overcome some of these deficiencies. Proximity-dependent labelling approaches are utilised by fusing a promiscuous mutant E. coli biotin ligase BirAR118G (BioID) to a protein of interest (bait), which allows biotinylation of neighboring and interacting proteins in the presence of biotin (Choi - Rhee et al., 2004; Cronan, 2005; Roux et al., 2012). Thus, biotinylated proteins are selectively captured by streptavidin-based affinity purification and subsequently identified by mass spectrometry. Because of strong covalent modifications and proteins’ labelling are executed in the native cellular environment, BioID-based proximity labelling tools enable the identification of weak interacting networks and allow the utilisation of insoluble protein compartments (Gupta et al., 2015; Kim et al., 2014; Li et al., 2017). However, the BioID labelling system is a slow kinetic process, which necessitates labelling with biotin for 18-24 h to produce sufficient biotinylated materials for proteomics analysis (Uezu et al., 2016; Youn et al., 2018).

Ascorbate peroxidase (APEX), a developed proximity labelling ligase in 2013, is characterised by a rapid labelling time (1 min or less) and subsequent generation of a snapshot of proximal proteins, and thus, it enables the identification even of components that interact transiently (Lam et al., 2015; Rhee et al., 2013). APEX has been successfully utilised in the characterization of the proteomes of the mitochondrial matrix, the primary cilium, and the ER-plasma membrane junctions, as well as in the composition profiling of the lipid droplet (Bersuker et al., 2018; Hung et al., 2017; Jing et al., 2015; Mick et al., 2015). However, utilisation of toxic H2O2 during labelling and high endogenous peroxidase activity limit APEX broad application on living samples and plants (Hung et al., 2016; Zhang et al., 2019).

More recently, Branon et al. directly evolved the *E. coli* biotin ligase BirA using yeast display and generated two promiscuous labelling variants, TurboID and miniTurbo, which enable sufficient proximity labelling in just 10 min with the use of non-toxic biotin (Branon et al., 2018). TurboID and miniTurbo have been demonstrated to probe different organellar proteomes in HEK cells (Branon et al., 2018). Because of the high labelling efficiency and lower temperature requirement, TurboID and miniTurbo have been utilised to profile interaction networks in *S. pombe*, *N. benthamiana*, and *Arabidopsis* (Larochelle et al., 2019; Mair et al., 2019; Zhang et al., 2019). Branon et al. also used the proximity labelling ability of TurboID and miniTurbo in *Drosophila* and *C. elegans* systems but without fusing any bait proteins to capture neighboring proteins (Branon et al., 2018).

Here, we applied TurboID-mediated proximity labelling in a wide variety of developmental stages and tissues in *Drosophila*. Using a cell-specific GAL4 driver, we also verified that TurboID can biotinylate the bait proteins, thus, making possible the identification of protein-protein interactions in individual cells. For the proof-of-principle, we used the TurboID labelling to analyse the proximate proteome of the cytoophidium, a newly identified filamentous structure containing the metabolic enzyme CTP synthase (CTPS). We show that by utilising TurboID-mediated biotinylation coupled with mass spectrometry, known CTPS-interacting proteins in *Drosophila* could be recovered. Our results suggest that TurboID is a feasible tool which can be used to characterise the proximate proteome of subcellular compartments in *Drosophila*.

## Results and Discussion

### TurboID-mediated protein biotinylation in *Drosophila*

The two promiscuous variants TurboID and miniTurbo, have been reported to biotinylate proteins in the presence of biotin in mammalian cells, *C. elegans*, *N. benthamiana* and *A. thaliana* (Branon et al., 2018; Mair et al., 2019). To map the proteome of CTPS cytoophidium in *Drosophila*, the initial concern we had to address was that the filamentous structures of CTPS should not be affected under recombinant expression with TurboID or miniTurbo (hereafter called TbID or miniTb).

Firstly, we checked for any conformational changes on CTPS cytoophidia by fusing TbID or miniTb containing V5 tag to the C-terminal of CTPS followed by transfection into *Drosophila* cultured S2 cells. Immunofluorescence results showed that CTPS retained its filamentous structures with TbID, while the conformation of CTPS tagged with miniTb became irregular and filaments were disrupted (Fig 2A). Branon et al. reported that miniTb is less stable than TbID likely due to removal of its N-terminal domain (Branon et al., 2018), which may explain that miniTb disrupted CTPS filamentous structures in *Drosophila* cultured cells. Considering the effects on CTPS cytoophidium conformation, we chose TbID as a promiscuous enzyme in our study.

**Fig. 1.**
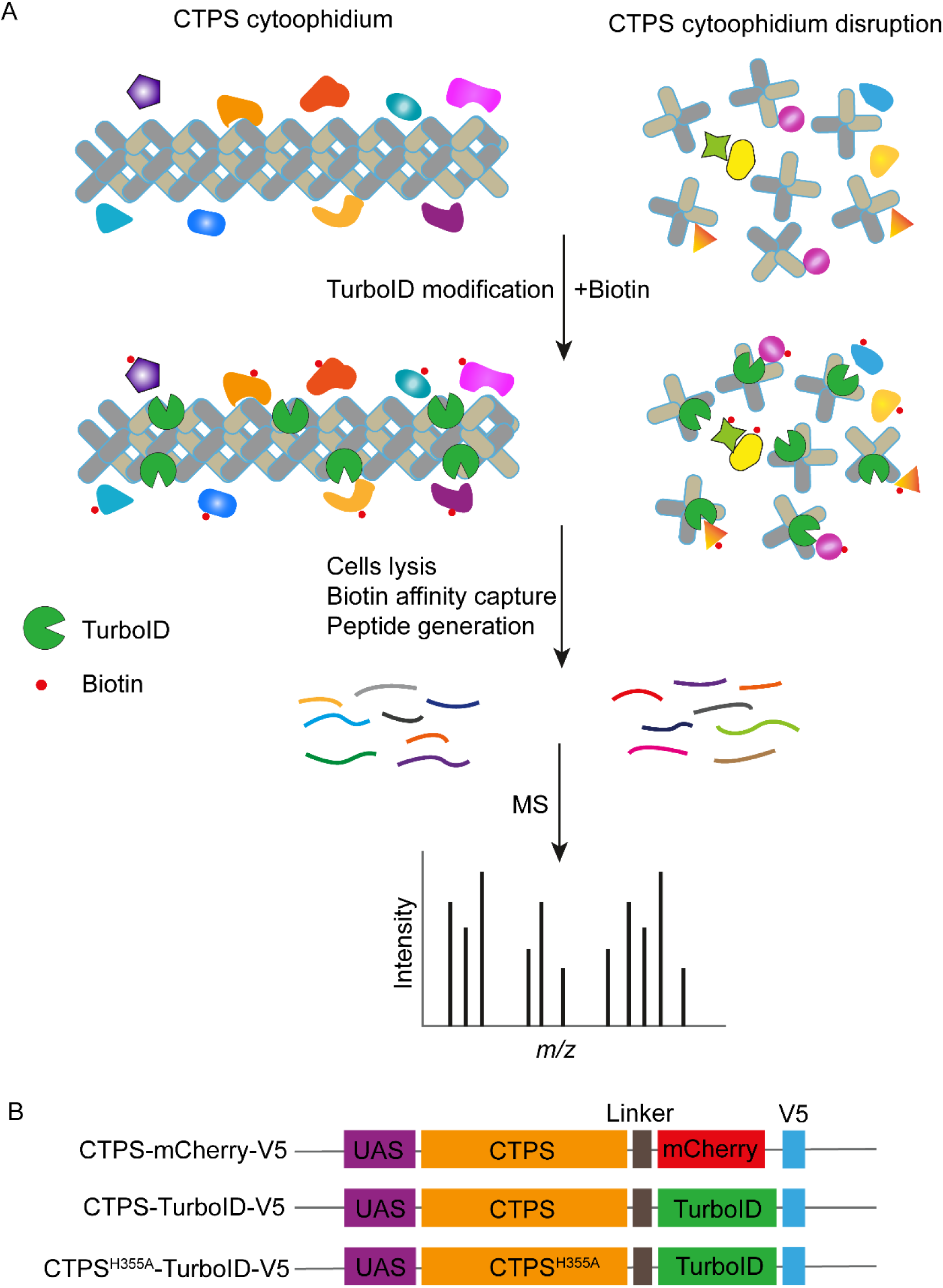
Model for TurboID-based labelling of CTPS cytoophidium proximate proteome. (A) TurboID was fused in-frame with wild type and mutant CTPS. Provided with biotin, TurboID can use biotin to biotinylate CTPS neighboring proteins. Cells are lysed and biotinylated proteins are captured using streptavidin beads. Subsequently, small peptides are generated by trypsin digestion and peptides are analyzed by mass spectrometry. Note that just a finite number of TurboID are shown in CTPS cytoophidium and disrupted cytoophidium. (B) Diagram of the expression cassettes used for the generation of transgenic flies. TurboID and V5 tag were fused to the C-terminal of wild type and mutant CTPS. mCherry was used as control. The same flexible linker was inserted into three cassettes.

**Fig. 2.**
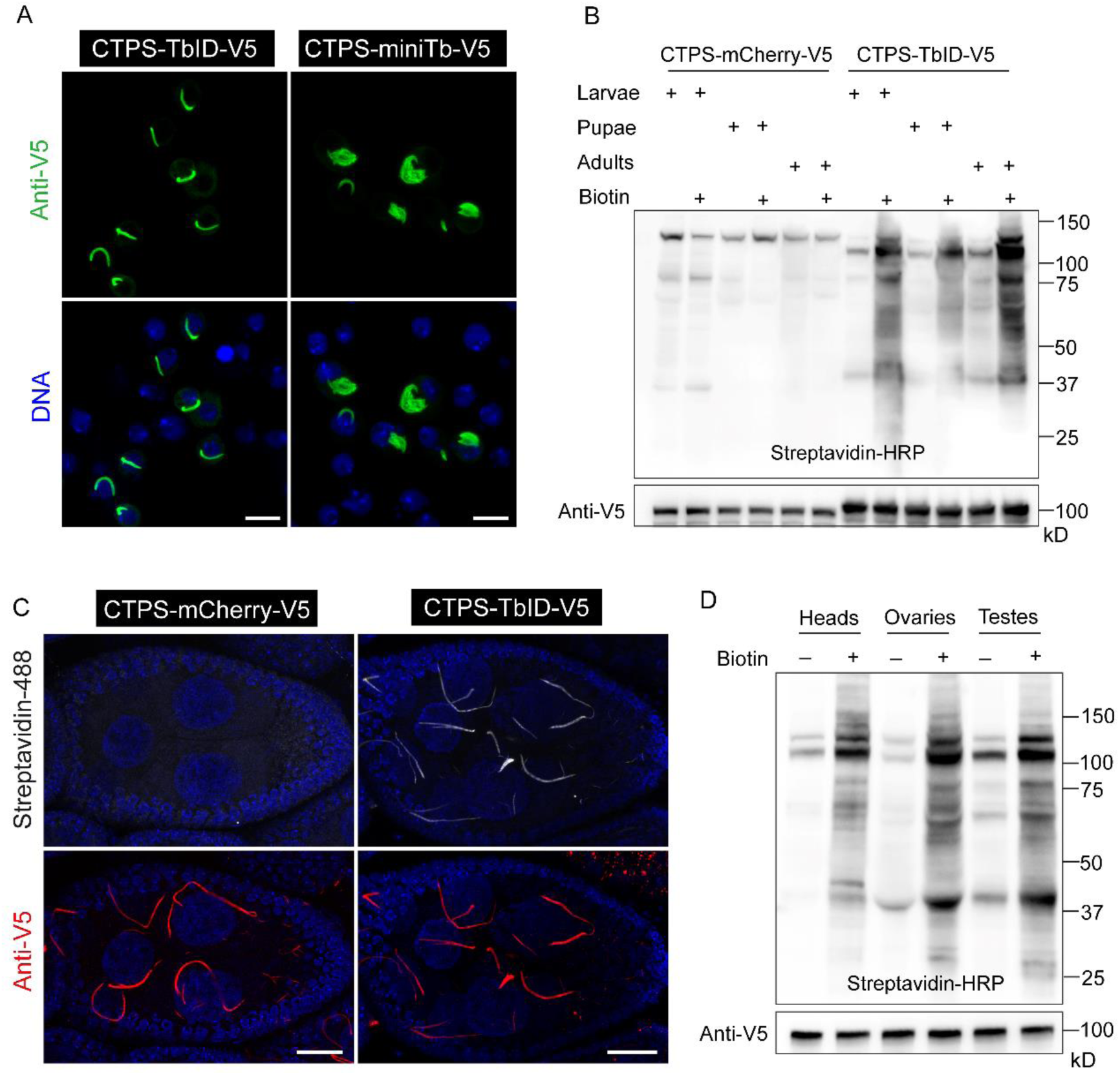
TurboID application in *Drosophila*. (A) CTPS-TbID and CTPS-miniTb containing V5 tag at C-terminal were cloned into pAc vectors and were transfected into *Drosophila* cultured S2 cells. Confocal images of cells are presented. DNA was labeled with Hoechst 33342 (blue). Scale bar, 10 μm. (B)Transgenic flies of UAS-CTPS-mCherry-V5 and UAS-CTPS-TbID-V5 were generated. Flies raised on either biotin-containing food or regular food from early embryo stages to larvae, pupae or adulthood, were collected and lysed and then blotted with Streptavidin-HRP to visualize biotinylated proteins. Anti-V5 antibody was used to detect the fused expression of CTPS-mCherry and CTPS-TbID, which was expressed ubiquitously via da-GAL4 driver. The molecular weight of CTPS-mCherry (98kD) is a little smaller than CTPS-TbID (105kD). (C) Both CTPS-mCherry and CTPS-TbID were specifically expressed in germline cells driven by nanos-GAL4. Ovaries from 14-day old flies grown on 100 μM biotin-containing food were dissected. Representative images of biotinylated proteins were obtained after detection by staining with streptavidin-488, while the expression of CTPS-mCherry and CTP-TbID was detected by anti-V5 blotting. Scale bar, 20 μm. (D) CTPS-TbID was expressed ubiquitously via da-GAL4 driver. Western blotting with streptavidin-HRP to visualize biotinylated proteins in different tissues from adult flies raised on either 100 μM biotin-containing food or regular food is presented. CTPS expression was detected by anti-V5 blotting.

To demonstrate TbID proximate labelling feasibility of CTPS in *Drosophila*, we initially generated UAS-CTPS-TbID and UAS-CTPS-mCherry transgenic flies, containing V5 tag at C-terminal of CTPS;, here, UAS-CTPS-mCherry served as a control group (Fig 1B). We then examined TbID wider application in all growth stages in *Drosophila*, and, to do so, we used da-GAL4 to drive CTPS-TbID and CTPS-mCherry ubiquitous expression. Flies, raised on either biotin-containing food or regular food from early embryo stages to larvae, pupae or adulthood, were collected and lysed. Streptavidin-HRP blotting results indicated that TbID biotinylated proteins in a wide variety of developmental stages in the presence of biotin, while no obvious labelling signals were detectable in any stages in CTPS-mCherry group (Fig 2B).

A previous study reported that CTPS cytoophidia exist in multiple tissues in *Drosophila*, such as brain, gut, testis, accessory gland, salivary gland and ovary (Liu, 2010). Based on that, we then dissected some tissues from adult flies in which CTPS-TbID and CTPS-mCherry were expressed via da-GAL4 driver. Western-blotting results using streptavidin-HRP revealed that TbID is extensively biotinylatilated in heads, ovaries, and testes, whereas no biotinylated proteins were detected in the control group (Fig 2D and Supplementary Fig. 1). Recently, Shinoda et al. knocked TbID into the C-terminal domain of caspase proteins’ gene loci by utilising CRISPR/Cas9 technology and labeled potential neighboring proteins in wings (Shinoda et al., 2019). In agreement with these studies, our results show that TbID can be used to label bait proteins in a wide range of developmental stages and tissues in *Drosophila*.

### TurboID labelling in *Drosophila* specific cells

After verifying the general applicability of TbID to proximity labelling in *Drosophila*, we wanted to further test whether TbID could biotinylate bait proteins and characterize local proteomes in individual cell types in *Drosophila*. A typical ovary in adult flies contains 16 ovarioles, each being tipped with the germarium which is followed by the growing egg chambers (Liu, 2010; Spradling, 1993). Each egg chamber includes three major cell types: one oocyte, 15 nurse cells and hundreds of follicle cells (Spradling, 1993). CTPS has been reported to form filaments in all three major cell types in ovaries (Liu, 2010). UAS-Gal4 is the most commonly used system for ectopic expression in *Drosophila*, and it can control the direct expression of genes of interest spatially and temporally, resulting in the desired expression pattern (White-Cooper, 2012).

Here we used nanos-GAL4, a driver controlling gene expression in germline stem cells and spermatogonia, to achieve CTPS direct expression in nurse cells and oocytes, but not in follicle cells in ovaries. We then dissected ovaries from 14-day old flies grown on biotin-containing food. Using anti-V5 antibody, we found that CTPS-mCherry and CTPS-TbID were both expressed and formed long and curved filamentous structures in nurse cells, whereas they did not in surrounding follicle cells, as expected (Fig 2C). The morphology of CTPS cytoophidia in nurse cells was similar to previous studies (Gitai et al., 2013).

In CTPS-mCherry group, no obvious labelling signals were detected after staining with streptavidin-AlexaFluor488 (Fig 2C). In contrast, in the case of CTPS-TbID, immunofluorescence images revealed that the biotinylated proteins are characterised by extensive signals and form patterns almost identical to CTPS cytoophidia (Fig 2C), indicating that the proteins in the vicinity of CTPS were preferentially biotinylated by TbID. Thus, our study shows that the TbID-based proximity system can be successfully used in labelling proteins of interest in desired cells using a cell-type-specific driver in *Drosophila*.

### Biotinylation of cytoophidia enabled by TurboID

CTP synthase is an essential enzyme that catalyzes the *de novo* formation of CTP from UTP and ATP using glutamate as nitrogen source (Levitzki et al., 1971). The enzyme consists of an N-terminal synthetase/amidoligase (ALase) domain that binds ATP and UTP, a helical interdomain linker and a C-terminal glutamine aminotransferase (GATase) domain (Endrizzi et al., 2004; Goto et al., 2004). The purified human CTP synthase 1 (hCTPS1) has been reported to polymerize to form filaments in the presence of UTP and ATP substrates, but not in the presence of CTP and ADP products, and filamentous structures of hCTPS1 have been shown to be formed by stacked tetramers (Lynch et al., 2017). A single histidine mutation on the tetramerization interface, H355A, renders hCTPS1 unable to polymerize into filaments in the presence of substrates (Lynch et al., 2017). Sun et al. have confirmed that the mutation H355A on mouse CTP synthase 1 (mCTPS1) also disrupts mCTPS1 assembly in mammalian cells (Sun and Liu, 2019a).

In order to characterize the proteome of CTPS cytoophidium, we worked in parallel with two distinct groups: CTPS formed filamentous structures in one group, while CTPS cytoophidia were disrupted in the control group. We aligned the protein sequences among *drosophila* CTPS (dCTPS), hCTPS1 and mCTPS1, and found that the identity of dCTPS to hCTPS1 and mCTPS1 was 65.66% and 66.16%, respectively (Supplementary Fig. 2). The alignment result showed that the amino acid histidine 355 (H355) was conserved among hCTPS1, mCTPS1 and dCTPS (Fig 3A).

**Fig. 3.**
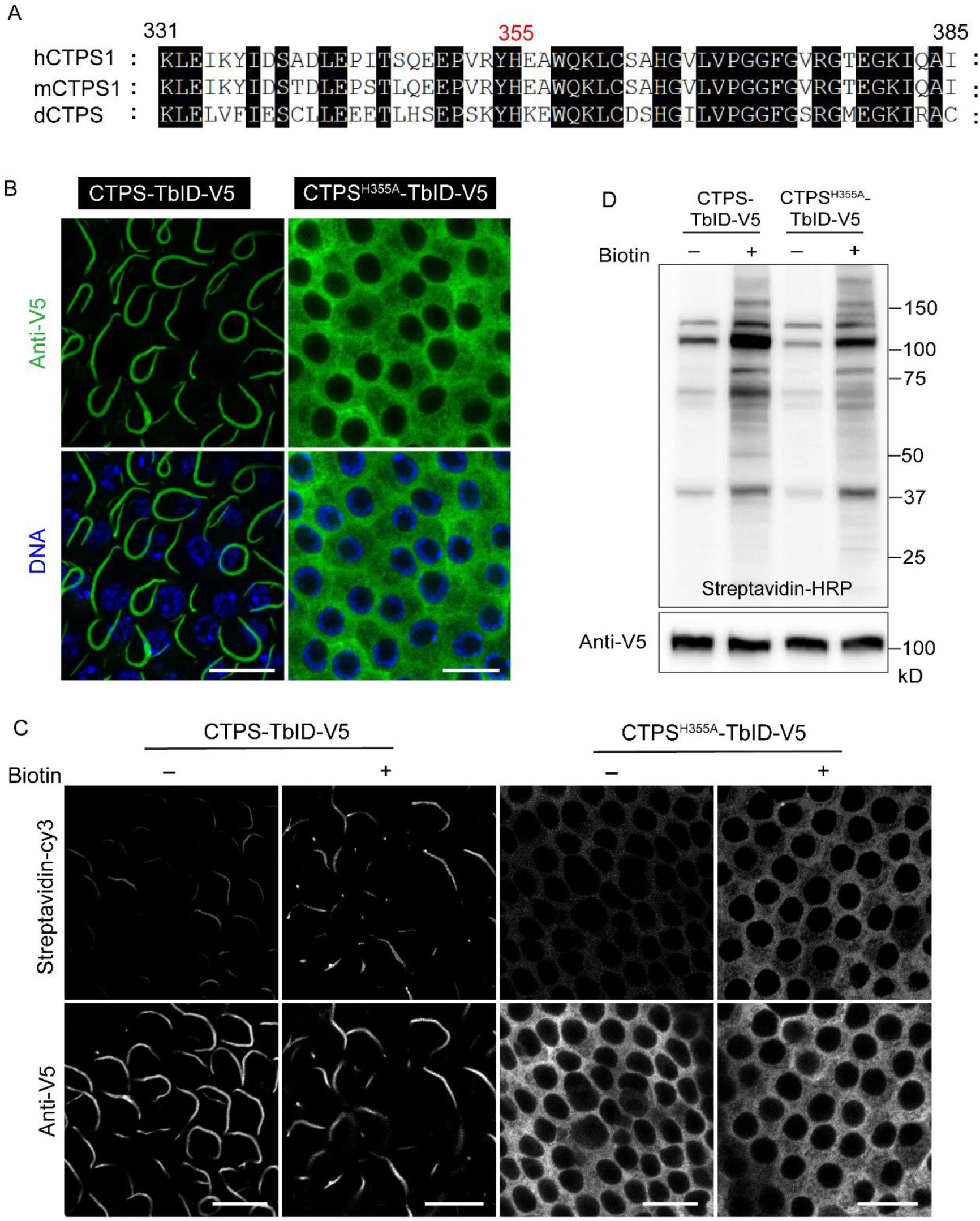
TurboID-mediated proximal biotinylation of CTPS cytoophidium and mutant CTPS. (A) Amino acid sequence alignment among hCTPS, mCTPS, and dCTPS. A partial view is presented here. (B) Immunostaining results of CTPS-TbID and CTPS^H355A^-TbID is shown in follicle cells. Scale bar, 10μm. (C) Ovaries were dissected from 14-day old flies and raised on either 100 μM biotin-containing food or regular food. Confocal images of labeled proteins detected by staining with streptavidin-Cy3 are presented, along with the expression of CTPS-TbID and CTPS^H355A^-TbID detected by anti-V5 blotting. All images were acquired from follicle cells. Scale bar, 10μm. (D) Streptavidin-HRP was used for the detection of labeled proteins, while anti-V5 antibody was used to detect the expression of CTPS-TbID and CTPS^H355A^–TbID. All ovaries samples were collected from 14-day old flies and CTPS-TbID and CTPS^H355A^–TbID were expressed using da-GAL4 driver in (B-D).

To detect whether H355A mutation impedes CTPS cytoophidium assembly in *Drosophila*, we generated a UAS-CTPS^H355A^-TbID transgenic fly by fusing a V5 tag at its C-terminal (Fig 1A). CTPS-TbID and CTPS^H355A^-TbID were both expressed by da-GAL4 driver, and ovaries from adult flies were used to examine the conformation of CTPS. Immunofluorescence results revealed that wild type CTPS formed long and curved filaments in follicle cells, while mutant CTPS showed a completely diffused distribution in cells (Fig 3B). These results indicated that H355A mutant disrupts CTPS polymerization in *Drosophila* follicle cells, providing an ideal control to characterize the neighboring proteome for CTPS cytoophidium. In order to reduce the background and contamination generated during sample preparation for mass spectrometry (MS) analysis, and considering that ovary serves as a classical research tool in *Drosophila*, we decided to use ovaries as models to identify the proteome of CTPS cytoophidium, rather than using whole flies.

Next, we explored the biotinylation of normal and disrupted CTPS cytoophidia mediated by TbID in ovaries. We expressed CTPS-TbID and CTPS^H355A^-TbID via da-GAL4 driver and dissected adult flies, grown on either biotin-containing food or regular food since the early embryo stages. V5 Western blotting results revealed a normal expression for the wild type and mutant CTPS in follicle cells (Fig 3C). Biotinylated proteins were detected using streptavidin-cy3 and immunofluorescence analysis showed extensive labelling signals for TbID, in the presence of biotin. Furthermore, we found that the biotinylated proteins had similar distribution patterns to the wild type, or disrupted cytoophidia, which suggested that the proximate proteins of CTPS-TbID/CTPS^H355A^-TbID were easily biotinylated (Fig 3C). In the absence of exogenous biotin, some weak labelling signals were also detected by streptavidin-cy3 (Fig 3C), which was expected because TbID has great efficiency and consumes endogenous biotin in cells in order to function, as was reported in previous studies (Branon et al., 2018; Larochelle et al., 2019). Additionally, using streptavidin-HRP blotting, we further confirmed that the neighboring proteins of CTPS and CTPS^H355A^ were biotinylatedby TbID (Fig 3D).

### TurboID-mediated CTPS cytoophidium proximate proteome

To characterize the proteome of CTPS cytoophidium dependent on TbID biotinylation in *Drosophila*, we expressed CTPS-TbID and CTPS^H355A^-TbID ubiquitously by da-GAL4 driver. Then, about 60 ovaries from 14-day old flies raised on biotin-containing food were collected, ground, and lysed. Biotinylated proteins were captured with streptavidin beads and subjected to on-bead trypsin digestion to generate peptides for analysis by mass spectrometry (MS). The three biological replicates for each group demonstrated good reproducibility, and showed a good correlation (Fig 4A).

**Fig. 4.**
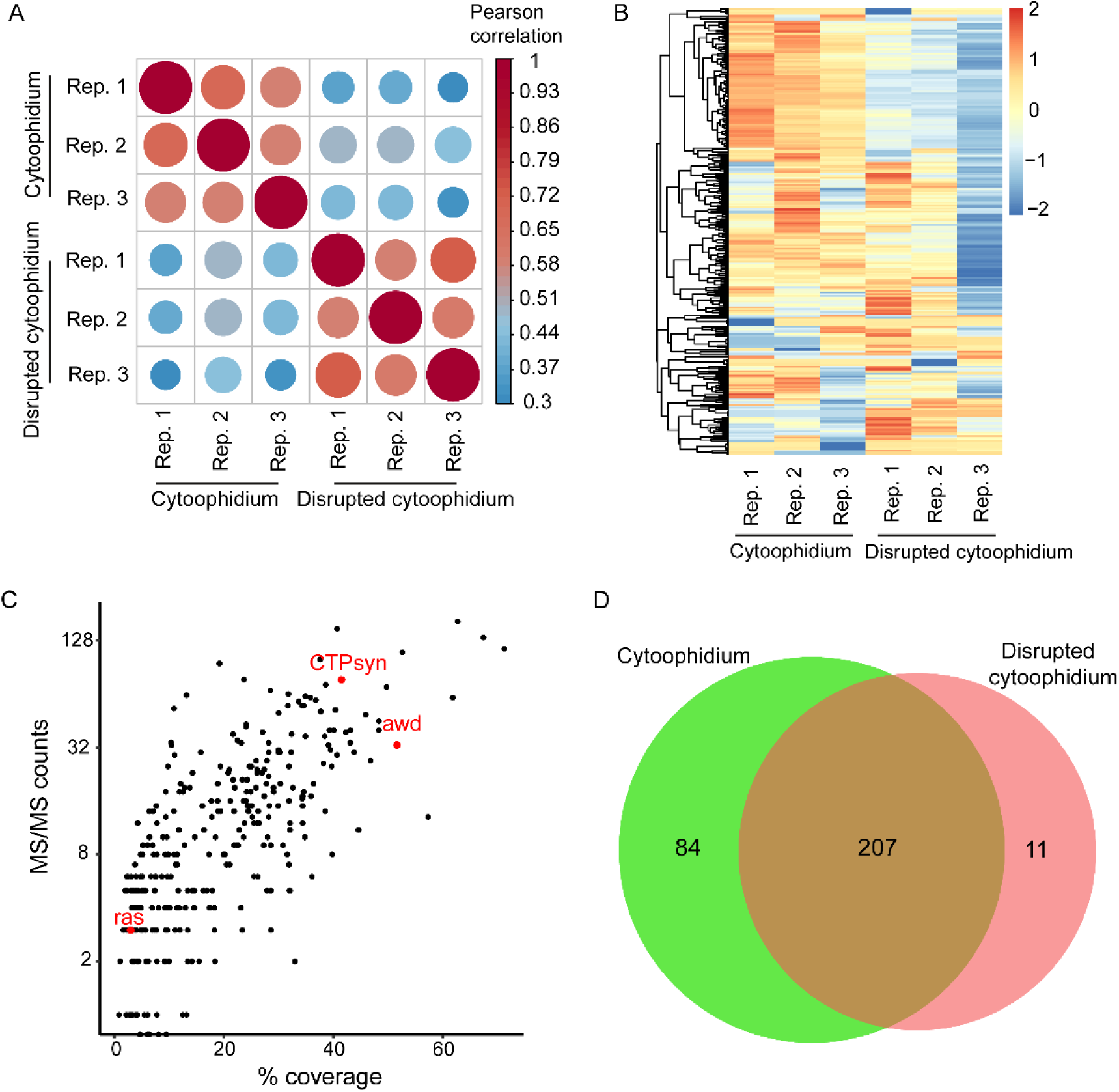
Characterization of proximate proteome of CTPS cytoophidium in *Drosophila*. CTPS-TbID and CTPS^H355A^-TbID were expressed by da-GAL4 driver. About 60 ovaries from adult flies grown on biotin-containing food were collected and prepared for mass spectrometry assay. Each experiment was repeated three times. (A) Pearson correlation coefficients between replicate (Rep.) mass spectrometry for cytoophidium and disrupted cytoophidium groups. (B) Hierarchical clustering of proteome of CTPS cytoophidium and disrupted cytoophidium in three replicates. (C) Scatter plots by MS/MS counts up the y-axis and percentage sequence coverage (amino acids) on the x-axis. Points corresponding to previously established CTPS interacting proteins in STRING database are labeled in red. (D) Venn diagram showing overlap and unique enriched proteins adjacent to CTPS cytoophidium and disrupted cytoophidium.

Then, we analyzed the biotinylated proteins adjacent to CTPS-TbID or CTPS^H355A^-TbID by hierarchical clustering, and our results revealed the differences among the proteome between normal CTPS cytoophidium and disrupted cytoophidium groups (Fig 4B). To assess the relative abundance of the characterized proteins, we analyzed MS/MS counts plotted against the protein sequence coverage (percentage of amino acid of a protein characterized by MS) by calculating the counts of all peptides matching to a specific protein (Larochelle et al., 2019; Schwanhausser et al., 2011). In addition to CTP synthase, another two known CTPS-interacting proteins (awd, ras) were found in our assay, and, as expected, CTPsyn was determined as a top hit (Fig 4C).

By differential expression analysis of the biotinylated proteins in two groups, we found that there were 207 proteins which overlapped, while 84 proteins were enriched in the vicinity of cytoophidium location. However, 11 proteins resided adjacently to the disrupted cytoophidium (Fig 4D, Supplementary Fig. 3). The enrichment of each protein, ranked by their fold change, is presented in Fig. 5A. We found that the subunits eIF-3p66, eIF3-S8, eIF3-S10, eIF-3p40 of the eukaryotic translation initiation factor 3 (eIF3), are enriched in the CTPS cytoophidium vicinity. Subunits of eIF2/2B complexes have been found to form filamentous structures in budding yeast (Noree et al., 2010; Shen et al., 2016). Our results raised the question of whether the subunits of eIF3, as the orthologs of eIF2/2B, form filamentous structures in *Drosophila* or whether they affect the assembly of CTPS cytoophidium. This gives us a direction towards a comprehensive investigation of intracellular compartments of metabolic enzymes and their proximate proteome.

**Fig. 5.**
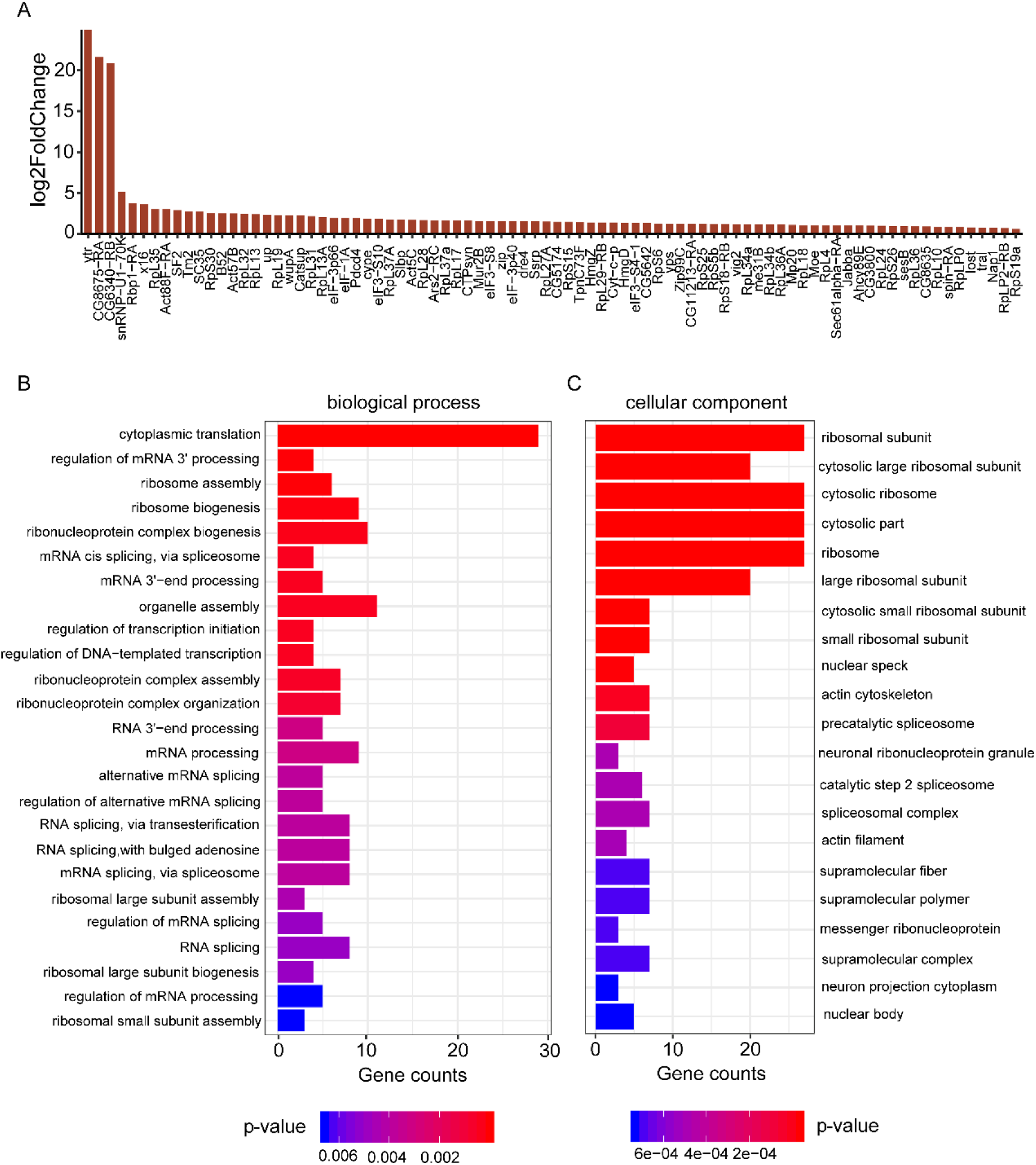
GO analysis of CTPS cytoophidium proximate proteome. (A) Bar blot list of 84 enriched neighboring proteins of CTPS cytoophidium compared to disrupted cytoophidium. (B) Enriched proximate proteins of CTPS cytoophidium classification based on biological processes. The single-item enrichment of p-value lower than 0.01 is shown and ranked by the p-value. (C) Enriched proximate proteins class distribution based on cellular components. p-value lower than 0.001 is shown.

Subsequently, we further categorized the enriched proteins of CTPS cytoophidium after performing gene ontology (GO) analysis. Several groups were overrepresented based on their biological process, including cytoplasmic translation, ribosome assembly, organelle assembly, ribonucleoprotein complex assembly, and ribosomal large subunit assembly (Fig 5B). Additionally, GO analysis revealed significant enrichment of cellular components related to the ribosomal subunit, cytosolic part, actin cytoskeleton, actin filament, supramolecular fiber, supramolecular polymer and supramolecular complex (Fig 5C). A previous study showed that CTPS functionally interacts with the intermediate filament, crescentin (CreS) to regulate cellular curvature in bacteria (Ingerson-Mahar et al., 2010). Here we identified some proteins, such as Act57B and Act5C, which serve as components of supramolecular fiber/polymer or polymeric cytoskeletal fiber. Whether Act57B and Act5C are involved in the synthesis of CTPS cytoophidium or they cooperate with CTPS to regulate cellular homeostasis has never been shown in *Drosophila*. Furthermore, enriched molecular functions included major clusters, such as the ones related to mRNA binding, actin binding, and cytoskeletal protein binding (Supplementary Fig. 4). In our study, we characterized the proteome of CTPS cytoophidium, providing a reference for future exploration of potential cellular functions of CTPS compartmentation, coordinated with its neighboring proteins.

Here, we demonstrate the feasibility of TurboID-mediated labelling in a wide range of developmental stages, tissues and specific cells in *Drosophila*. We utilised TurboID-mediated labelling, as a proof-of-principle, for studying the proteome of CTPS cytoophidia. Our study describes a feasible approach to explore the proximate proteins and cellular functions of subcellular compartments of metabolic enzymes in vivo.

## Materials and Methods

### Construction of plasmids

*Drosophila* codon optimized TurboID and miniTurbo sequences were synthesized. CTPS, TurboID and miniTurbo containing V5 epitope were amplified by PCR (Vazyme, Cat. # P505-d3). pAc 5.1 vector was digested by EcoR I and Not I (NEB). The amplified CTPS, TurboID-V5, and miniTurbo-V5 were inserted into pAc vector by seamless cloning (Vazyme, Cat. # C113-02) to result in plasmids pAc 5.1 CTPS-TurboID-V5 and pAc 5.1 CTPS-miniTurbo-V5. To induce H355A directed point mutation in CTPS, the following primers were used to amplify wild type CTPS encoding sequence: F: 5’-GAGCAAGTACGCCAAGGAGTGGCAGAAGCTATGCGATAGCCA-3’ R: 5’-CACTCCTTGGCGTACTTGCTCGGCTCAGAATGCAAAGTTTCC-3’ To create pUAS-CTPS-mCherry-V5, pUAS-CTPS-TurboID-V5 and pUAS-CTPS^H355A^-TurboID-V5 plasmids, mCherry-V5, TurboID-V5, and CTPS^H355A^-V5 were amplified and inserted into pUAS vector by seamless cloning. pUAS vector was digested using Not I and Kpn I (NEB).

### Cell culture

S2 cell line was maintained in Schneider’s *Drosophila* medium (Gibco) supplemented with 10% fetal bovine serum (FBS) at 28°C incubator. Cell transfections were carried out using Effectene Transfection Reagent (QIAGEN) according to manufacturer’s instructions.

### Transgenic flies and *Drosophila* culture

Three transgenic fly lines were established in our study. To express ligase TurboID ubiquitously in flies, transgenic flies were crossed with da-GAL4 or nanos-GAL4 driver flies and recombinants were generated. All flies were raised at 25°C on either standard cornmeal food or 100 μM biotin-containing food accordingly.

### Immunofluorescence

For S2 cells, after 36-hour transfection, cells cultured on glass slides were fixed with 4% (v/v) paraformaldehyde in PBS for 20 min. Cells were washed three times with PBS and then permeabilized with 0.2% Triton X-100 for 15 min. After blocking with 5% (w/v) bovine serum albumin (BSA) in PBS for 1 hour, primary antibody incubation with anti-V5 antibody (1:500, Invitrogen, Cat. # 460705) in PBS was carried out overnight at 4°C. Following three washes in PBS, cells were incubated with Alexa Fluor 488-labeled secondary antibody (1:500, Invitrogen, Cat. # A11029) and Hoechst 33342 (1:10000, Bio-Rad, Cat. # 151304) for 1 hour.

For *Drosophila*, ovaries from 14-day flies were dissected in Graces Insect Med Sup (Life) and then fixed with 4% (v/v) paraformaldehyde in PBS for 20 min. Ovaries were incubated with anti-V5 antibody (Invitrogen) in PBS containing 0.3% Triton X-100 and 0.5% horse serum overnight. To detect the distribution of biotinylated proteins and expression of ligase, ovaries were incubated with Alexa Fluor 488-labeled secondary antibody (1:500, Thermo Fisher) and Streptavidin-Cy3 (1:300, Jackson, Cat. # 016-160-084) containing Hoechst 33342 (Bio-Rad) overnight before confocal imaging.

### Protein Extraction and Western blotting

To extract proteins from flies of different developmental stages, larvae, pupae, and adult flies were collected and frozen with liquid nitrogen for 1 min. For protein extraction from different tissues, the tissues were dissected in Graces Insect Med Sup (Life) and then were frozen with liquid nitrogen for 1 min. Samples were prepared with RIPA lysis buffer and 1X protease inhibitor cocktail (Bimake) and then ground for 10 min. Following 20 min incubation on ice, samples were centrifuged at 13,000 rpm at 4 °C for 30 min. The supernatants were collected and boiled with 1X protein loading buffer at 95 °C for 10 min. Lysates were separated by 4-20% SDS-PAGE gels, followed by transferring to PVDF membranes (Roche). After blocking with 5% BSA in TBST for 1 hour, membranes were incubated with HRP-conjugated antibody (1:3000, Cell Signaling, Cat. # 3999s) for 1hour. After three times washing in TBST, biotinylated proteins were visualized using enhanced chemiluminescence system. To detect the recombination expression of ligase, blocked membranes were incubated with anti-V5 antibody (1:3000, Invitrogen) at 4 °C overnight. Following three washes in TBST, membranes were incubated with HRP-linked anti-mouse antibody (1:3000, Cell Signaling, Cat. # 7076s) for 1 hour. Then,membranes were washed with TBST three times, before ligase expression was visualised by enhanced chemiluminescence system.

### Mass spectrometry sample preparation

About 60 ovaries from 14-day old adult flies grown on 100 μM biotin-containing food were dissected, then fixed with 4% (v/v) paraformaldehyde in PBS for 20 min, followed by washing with cold PBS one time, and then incubated with 1 ml lysis buffer at 4 °C. After shaking for 1 h, the lysate was spun down at 4 °C for 10 min. The supernatant was transferred into new tubes, with the addition of urea and DTT to a final concentration of 8 M and 10 mM. The lysate was incubated at 56 °C for 1 hour, then treated with 25 mM iodoacetamide in the dark for 45 min to aminocarbonyl modify the Cys site of proteins. 25 mM DTT was added to terminate the modification. 50 μL Streptavidin Magnetic Beads (NEB, Cat. # S1420S) were washed with 500 μL PBS three times and then resuspended into the lysate. Subsequently, the lysate along with with 50 μL beads was incubated on a rotator at 4°C overnight. The beads were washed with the following buffers: twice with buffer 1 (50 mM Tris8.0, 8 M urea, 200 mM NaCl, 0.2% SDS), once with buffer 2 (50 mM Tris8.0, 200 mM NaCl, 8 M urea), twice with buffer 3 (50 mM Tris8.0, 0.5 mM EDTA, 1 mM DTT), three times with buffer 4 (100 mM ammonium carboxylate), and finally the beads were resuspended in 100 μL buffer 4. Trypsin, 4 μg (Promega, Cat. # v5113) was added to digest proteins to generate peptides overnight at 37 °C. The peptides were collected with ziptip by the addition of 1% formic acid, then washed with 0.1% TFA (Sigma, Cat. # 14264) and eluted in 50 μl of 70% ACN (Merck Chemicals, Cat. # 100030)-0.1% TFA. The peptides were analyzed on an Orbitrap Fusion.

### Mass spectrometry data analysis

The UniProt *Drosophila* melanogaster protein database (Proteome ID: UP000000803), and database for proteomics contaminants from MaxQuant were used for database searches. Reversed database searches were used to evaluate the false discovery rate (FDR) of peptide and protein identifications. Two missed cleavage sites of trypsin were allowed. The FDR of both peptide characterization and protein characterization was set to be 1%. The options of “Second peptides”, “Match between runs”, and “Dependent peptides” were enabled. For differential expression analysis, the limma-based approach evolutionarily in R 3.6.1 was used. The data were log2 transformed and centered, and the statistical significance of the biological repeats of CTPS cytoophidium and disrupted cytoophidium control was tested using a modified t-test in limma 3.40.0. Enriched proteins of CTPS cytoophidium with fold change > 1.5 and p-value < 0.05 were defined as up-regulated proteins and those with fold change < 0.67 and p-value < 0.05 were defined as down-regulated proteins. Functional enrichment analysis was followed to define the enriched proteins of CTPS cytoophidium. clusterProfiler software was used to obtain enriched GO terms corresponding to biological processes (BP), cellular components (CC), and molecular functions (MF) (Yu et al., 2012). Only those categories with p-adjust lower than 0.05 were considered to be reliable. Previously reported CTPS-interacting proteins were obtained from STRING database (https://string-db.org).

## Acknowledgments

This work was supported by ShanghaiTech University. We thank Tiezhu Shi for the bioinformatics analysis and data visualization, Chengqian Zhang for the assistance with the mass spectrometry experiments and Xiaoming Li for providing the training on confocal microscopy. We are grateful to the Molecular Imaging Core Facility and the Mass Spectrometry Facility at School of Life Science and Technology, ShanghaiTech University for providing the necessary technical support. We thank Christos Andreadis for reading the manuscript.

## Author Contributions

B.Z. carried out all the experiments with the help of Y.Z. B.Z. conducted all the data analyses with the help of all the authors. J.L. supervised all experiments and data analyses and supervised the project. B.Z. and J.L. wrote the manuscript.

## Supplementary Information

**Supplementary Fig. 1.**
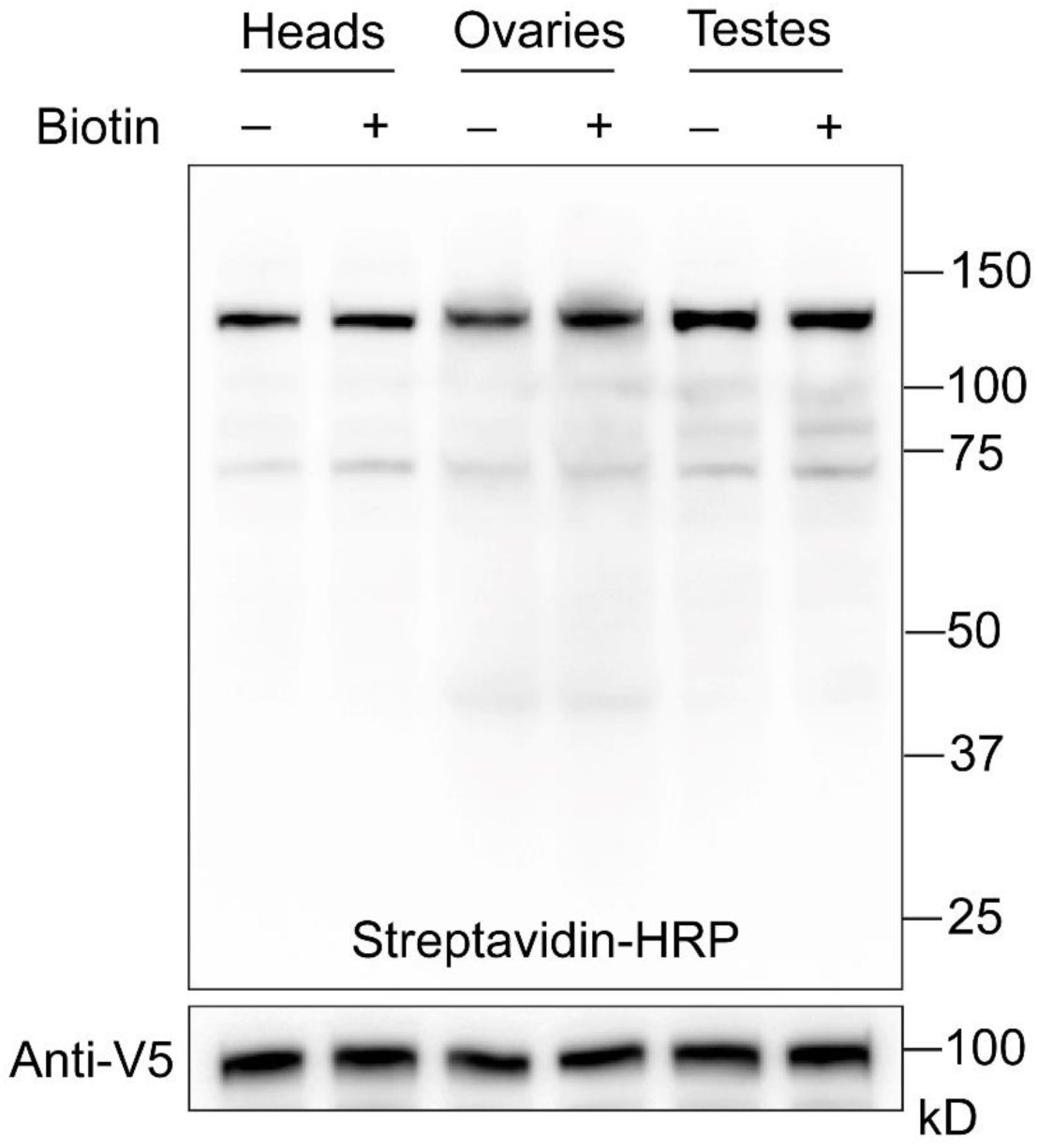
CTPS-mCherry was expressed ubiquitously via da-GAL4 driver. Western blotting visualizing biotinylated proteins in different tissues from adult flies with streptavidin-HRP. CTPS-mCherry expression was detected by anti-V5 blotting.

**Supplementary Fig. 2.**
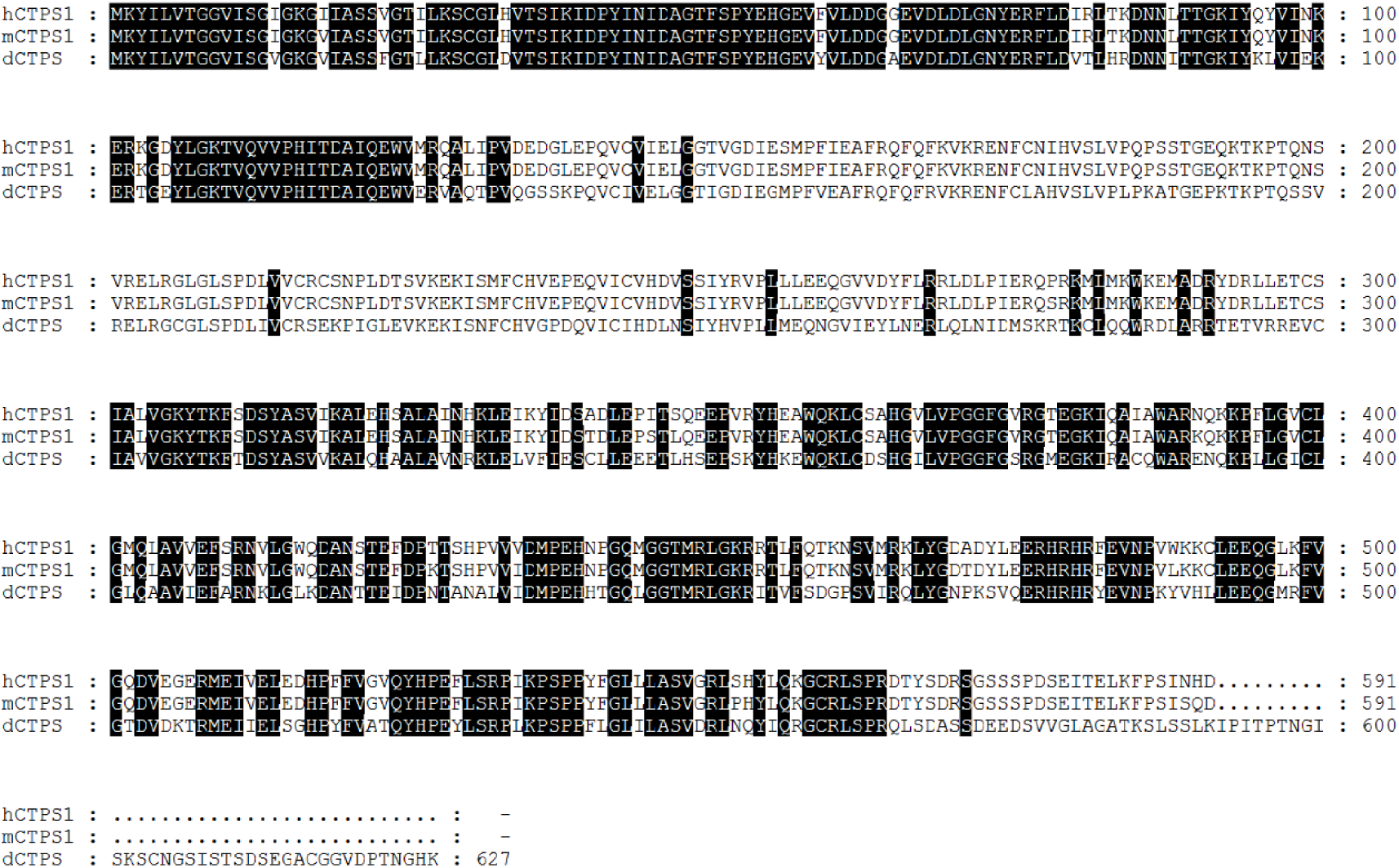
Sequence alignment result of hCTPS1, mCTPS1, and dCTPS. Conserved amino acids were indicated in black.

**Supplementary Fig. 3.**
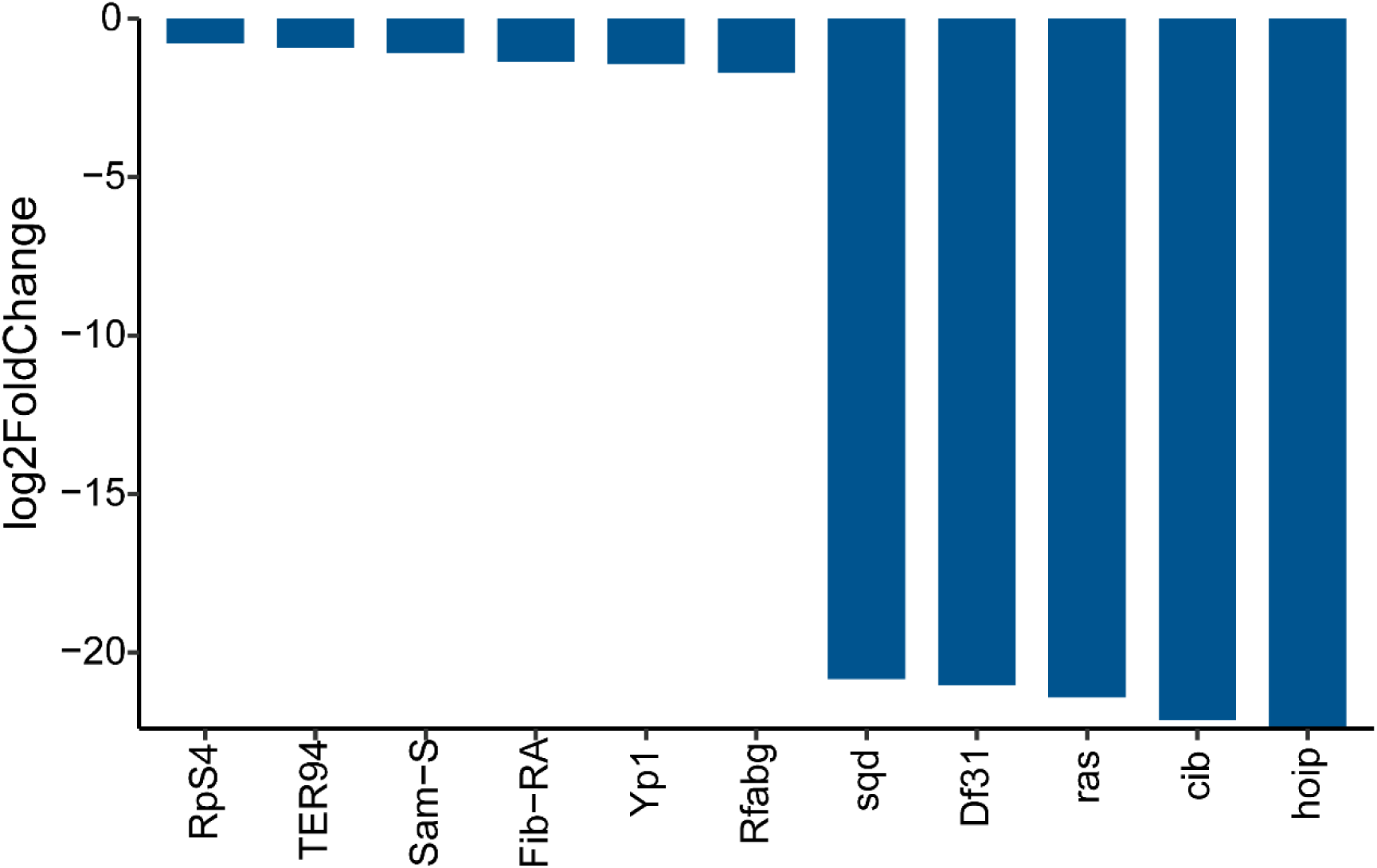
List of 11 enriched proteins residing adjacently to disrupted CTPS cytoophidium.

**Supplementary Fig. 4.**
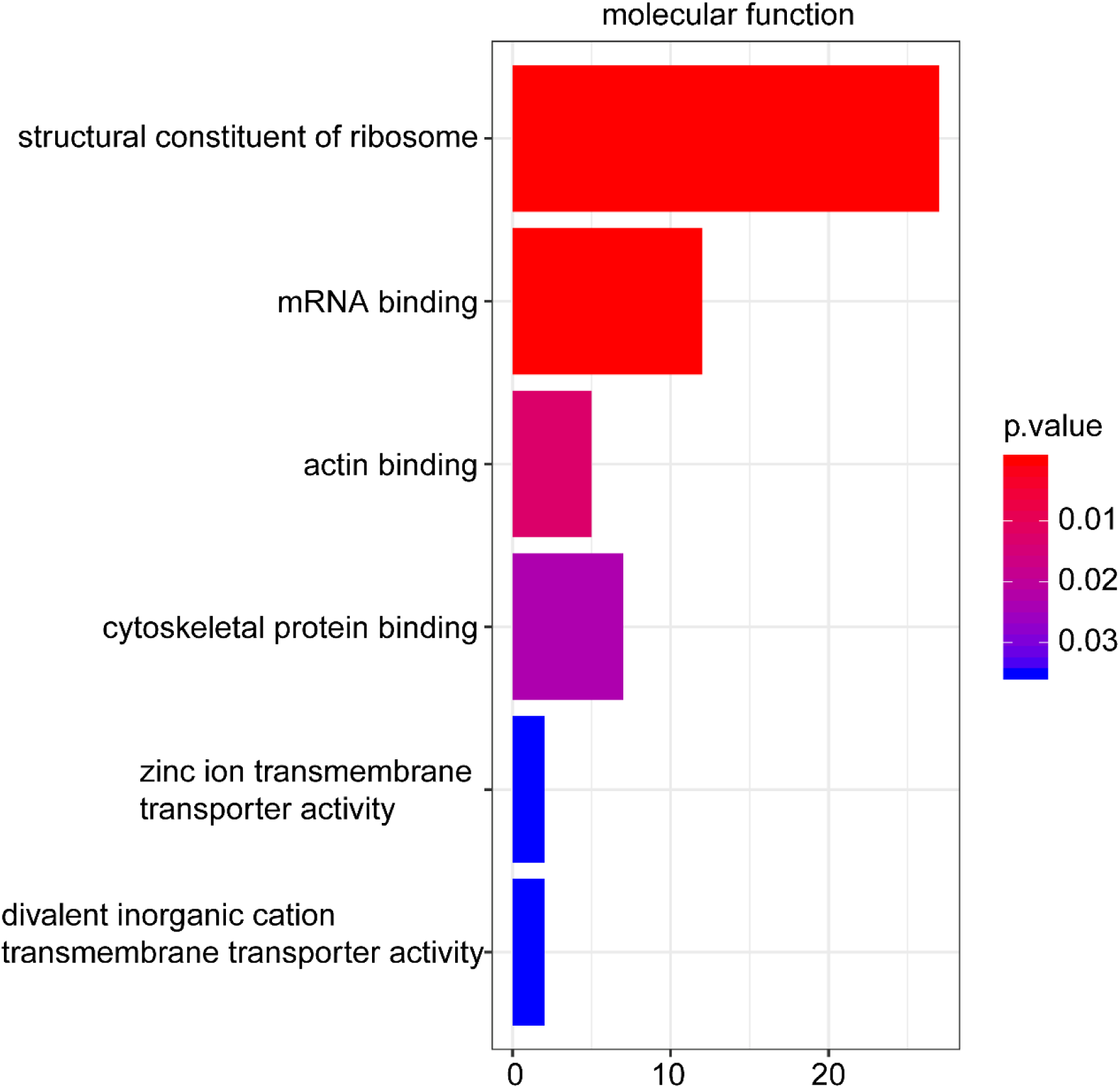
Enriched proximate proteins of CTPS cytoophidium classification based on molecular functions. The single-item enrichment of p-value lower 0.05 is shown and ranked by p-value.

